# A PI(3,5)P2/ESCRT-III axis terminates STING signalling by facilitating TSG101-mediated lysosomal microautophagy

**DOI:** 10.1101/2024.05.26.595979

**Authors:** Tsumugi Shoji, Ayumi Shinojima, Satoshi Kusumi, Daisuke Koga, Kojiro Mukai, Jun Nakayama, Shigeki Higashiyama, Yoshihiko Kuchitsu, Tomohiko Taguchi

## Abstract

Stimulator of interferon genes (STING) is critical for the type I interferon response to pathogen- or self-derived cytosolic DNA. STING is degraded by the endosomal sorting complexes required for transport (ESCRT)-driven lysosomal microautophagy (LMA), the impairment of which leads to sustained inflammatory responses. It has been unknown how ESCRT targets STING directly to lysosomes. Here, through kinase inhibitor screening and knockdown experiments of all the individual components of ESCRT, we show that STING degradation requires PIKfyve (a lipid kinase that generates PI(3,5)P_2_) and CHMP4B/C (components of ESCRT-III subcomplex). Knockdown of Pikfyve or Chmp4b/c results in the accumulation of STING vesicles of a recycling endosomal origin in the cytosol, leading to sustained type I interferon response. CHMP4B/C localize at lysosomes and their lysosomal localization is abolished by interference with PIKfyve activity. Our results identify lysosomal ESCRT-III as a PI(3,5)P_2_ effector, reveal a role of the less characterized phosphoinositide PI(3,5)P_2_ in lysosomal biology, and provide insights into the molecular framework that distinguishes LMA from other cellular processes engaged with ESCRT.

## Introduction

Stimulator of interferon genes (STING)^1^, also known as MITA^2^/ERIS^3^/MPYS ^4^/TMEM173, is an endoplasmic reticulum (ER)-localized transmembrane protein essential for control of infections of DNA viruses and tumor immune surveillance^5-7^. With binding to cyclic GMP-AMP^8^, which is produced by cyclic GMP-AMP synthase^9^ in the presence of cytosolic DNA, STING migrates to the trans-Golgi network (TGN) where STING triggers the type I interferon and proinflammatory responses through the activation of interferon regulatory factor 3 (IRF3) and nuclear factor-κB^10-12^. STING further migrates to lysosomes for its degradation through recycling endosomes (REs), so that the STING-signalling is terminated^13^. We have recently shown that STING degradation is executed by lysosomal microautophagy (LMA), in which the limiting membrane of lysosomes directly encapsulates degradative substrates in the cytosol^13, 14^. We further revealed that the recognition of K63-ubiquitinated STING by TSG101, a subunit of the endosomal sorting complexes required for transport (ESCRT), was required for STING degradation by LMA^13^.

ESCRT is the evolutionarily conserved protein complex that plays a crucial role in various cellular processes, including endosomal sorting of plasma membrane proteins to lysosomes, cytokinesis, and viral budding^15^. ESCRT in mammals consists of four subcomplexes, ESCRT-0 (HRS, STAM1, and STAM2), ESCRT-I (TSG101, VPS28, VPS37A/B/C/D, MVB12A/B, and UBAP1), ESCRT-II (VPS22, VPS25, and VPS36), and ESCRT-III (CHMP1A/B, CHMP2A/B, CHMP3, CHMP4B/C, CHMP5, CHMP6, CHMP7, and IST1)^16^. In the process of the endosomal sorting, HRS recognizes ubiquitinated transmembrane proteins on membrane, which is followed by the sequential recruitment of ESCRT-I, ESCRT-II, and ESCRT-III. Finally, Vps4 is recruited onto ESCRT-III and disassembles polymerized ESCRT, leading to the generation of intraluminal vesicles (ILVs) that are eventually subjected to lysosomal degradation^15, 16^. In contrast to the mechanistic insights into the ESCRT function on endosomes, the ones into the ESCRT function on lysosomes, have not yet been fully elucidated.

In this article, through kinase inhibitor screening and knockdown experiments of all the individual components of ESCRT, we sought to elucidate the molecular mechanism that distinguishes LMA from other cellular processes engaged with ESCRT.

## Results

### PIKfyve inhibitors suppress STING degradation

To identify molecules required for STING degradation, we sought to set up a high-throughput screening system with the Nano-Glo^®^ Dual-Luciferase^®^ Reporter. Firefly luciferase (FLuc) and Nanoluciferase (NLuc) were used for expression control and for quantitation of STING levels, respectively. The coding sequences of FLuc and NLuc-tagged Sting were linked by a P2A self-cleavage site for equimolar expression of both proteins (Fig. 1a). FLuc-P2A-NLuc-STING was stably expressed in Sting^-/-^ mouse embryonic fibroblasts (MEFs), and characterized for its competence in signalling activity and for its susceptibility to degradation (Fig. S1a-b).

**Fig. 1.**
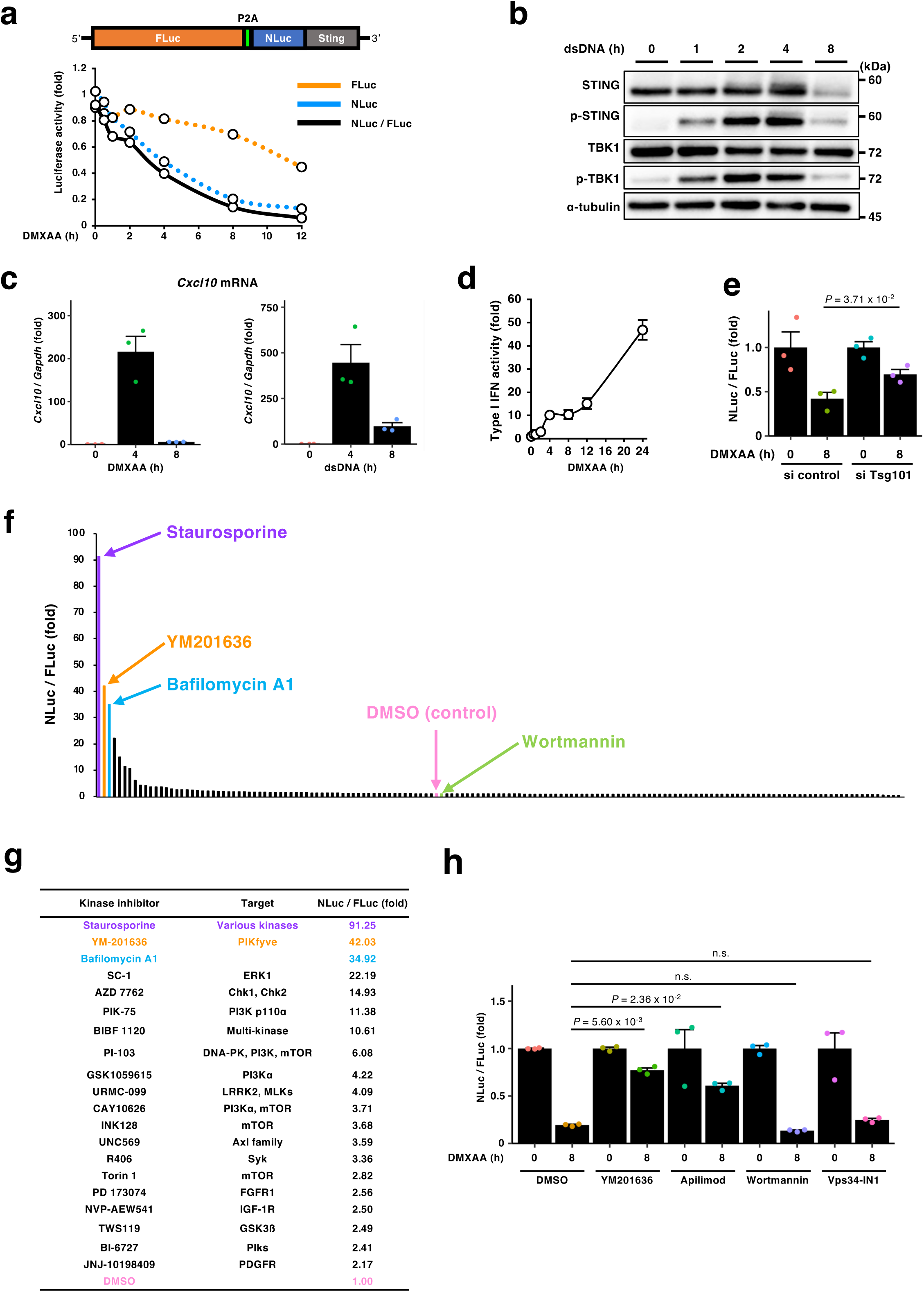
PIKfyve inhibitors suppress STING degradation. **a**, A schematic illustration of the plasmid encoding FLuc-P2A-NLuc-mouse Sting. Sting^-/-^ MEFs stably expressing FLuc-P2A-NLuc-STING were stimulated with DMXAA for the indicated times. The luminescence of FLuc and NLuc was quantitated at each time point, and the NLuc/FLuc ratio was plotted. **b-e**, The cells in (**a**) were stimulated with double-stranded DNA or DMXAA for the indicated times. Cell lysates were analyzed by western blot in (**b**). The expression of Cxcl10 was quantitated with qRT-PCR in (**c**). The activity of type I interferon (IFN) in the cell supernatants in (**d**). The effect of Tsg101 knockdown in (**e**). The cells treated with siRNA for 72 h were used. **f**, The cells in (**a**) were simultaneously treated with kinase inhibitors and DMXAA for 18 h. The NLuc/FLuc ratio from the cells treated with 1 µM inhibitors and DMXAA was then normalized to that from DMSO-treated cells. **g**, The kinase inhibitors ranked within the top 20 were listed. As positive control for the inhibition of STING degradation, the data with Baf. A1 were shown. **h**, The cells in (**a**) were treated with the indicated inhibitors (1 µM) for 2 h, and then stimulated with DMXAA for 8 h. Data are presented as mean ± standard error of the mean.

The cells were stimulated with DMXAA (a membrane-permeable STING agonist) for the indicated times, and the luciferase activities were measured. We found a time-dependent decrease of NLuc-STING (Fig. 1a), activation of TBK1 (Fig. 1b and Fig. S2 a), induction of Cxcl10 expression (Fig. 1c), and type I interferon production (Fig. 1d). In line with the previous findings that STING degradation requires STING translocation from the ER^17^, interference of the ER-Golgi traffic with brefeldin A (BFA) or with Sar1a/b-knockdown suppressed NLuc-STING degradation (Fig. S2b-d). Inhibition of vacuolar H^+^-ATPase with Bafilomycin A1 (Baf.A1) or with knockdown of Atp6v1b2, a component of subunit B of vacuolar ATPase, suppressed the degradation of NLuc-STING (Fig. S2c and e). Knockdown of Tsg101^13^, a gene essential for lysosomal microautophagic degradation of STING, suppressed the degradation of NLuc-STING (Fig. 1e).

154 kinase inhibitors (Table S1) were then examined for their ability to inhibit STING degradation. Cells were stimulated with DMXAA for 18 h in the presence of kinase inhibitors, and the NLuc/FLuc ratio was calculated. We found that 64 kinase inhibitors suppressed STING degradation more effectively than DMSO used as a vehicle (Fig. 1f-g). Among them, staurosporine and YM201636^18^ strongly inhibited STING degradation as Baf.A1. Staurosporine is a non-selective inhibitor of protein kinases, therefore we did not characterize it further. YM201636 is a selective inhibitor of PIKfyve^19^, a lipid kinase that generates phosphatidylinositol 3,5-bisphosphate [PI(3,5)P_2_] from phosphatidylinositol 3-phosphate [PI(3)P]. Another highly selective PIKfyve inhibitor apilimod^20^ also suppressed STING degradation, supporting a role of PIKfyve in STING degradation (Fig. 1h). Of note, wortmannin and Vps34-IN1, inhibitors of phosphatidylinositol 3-kinase (PI3K) that generates PI(3)P, did not inhibit STING degradation (Fig. 1f and 1h).

### PIKfyve is required for termination of STING signalling

We next examined a role of PIKfyve in termination of STING signalling. Immortalized MEFs (iMEFs) were stimulated with DMXAA in the presence of YM201636 or apilimod for the indicated times (Fig. 2a). In control cells, phosphorylated TBK1 (pTBK1) and phosphorylated STING (pSTING), the hallmarks of STING activation, emerged 2 h after stimulation, and then diminished. In contrast, in cells treated with YM201636 or apilimod, p-TBK1 and p-STING lingered up to 8 h after stimulation. The treatment with apilimod resulted in the increased expression of Cxcl10 (Fig. 2b). RNAseq analysis showed the upregulation of various proinflammatory genes with apilimod after 12-h stimulation (Fig. 2c: lane 3 versus lane 4). Knockdown of Pikfyve in primary MEFs retarded the disappearance of pTBK1 (Fig. 2d-e) and enhanced expression of Cxcl10 and Ccl5 (Fig. 2f). These results suggested that PIKfyve and its product PI(3,5)P_2_ were essential for the termination of STING signalling.

**Fig. 2.**
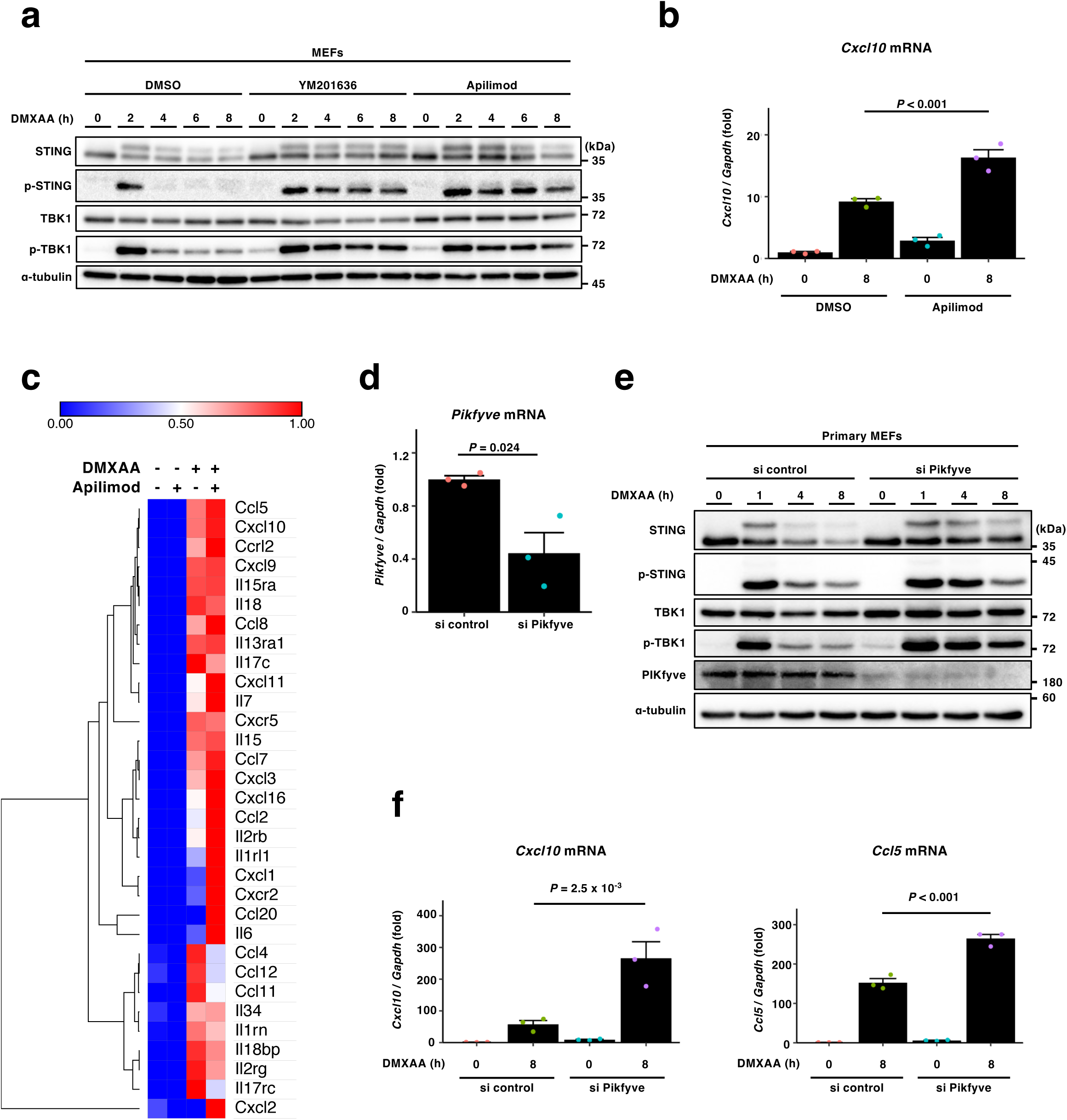
PIKfyve is required for STING degradation and termination of STING signalling. **a,** Cells were treated with DMXAA in the presence of PIKfyve inhibitors (1 µM) for the indicated times. Cell lysates were analyzed by western blot. **b,** Cells were treated with DMXAA in the presence of apilimod (1 µM) for 8 h. The expression of Cxcl10 was quantitated with qRT-PCR. **c,** A heatmap based on the expressional profiles of proinflammatory genes in RNA-seq analysis. Cells were pretreated with apilimod (1 µM) for 2 h, and stimulated with DMXAA for 12 h. The expression of genes was quantitated with RNAseq. **d,** Cells were treated with the indicated siRNAs for 96 h. The expression of Pikfyve was quantitated with qRT-PCR. **e**, Cells treated with the indicated siRNAs were stimulated with DMXAA for the indicated times. Cell lysates were analyzed by western blot. **f,** Cells treated with the indicated siRNAs were stimulated with DMXAA for 8 h. The expression of Cxcl10 and Ccl5 was quantitated with qRT-PCR. Data are presented as mean ± standard error of the mean.

### PIKfyve is required for LMA degradation of STING

The sustained STING signalling in Pikfyve-knockdown cells may be due to the impaired LMA degradation of STING. In this scenario, STING is expected to accumulate outside the lysosomes. To examine this, Sting^-/-^ MEFs that stably express mRuby3-STING and EGFP-tagged LAMP1 (a lysosomal protein) were prepared. The cells were stimulated with DMXAA for 12 h in the presence of lysosomal protease inhibitors, and examined by Airyscan imaging. In control cells, mRuby3-STING accumulated inside lysosomes (Fig. 3a). In contrast, mRuby3-STING was hardly observed in lysosomal lumen in apilimod-treated cells. These results suggested the defect of lysosomal encapsulation of STING. As reported, apilimod treatment resulted in dilation of lysosomes^21^.

**Fig. 3.**
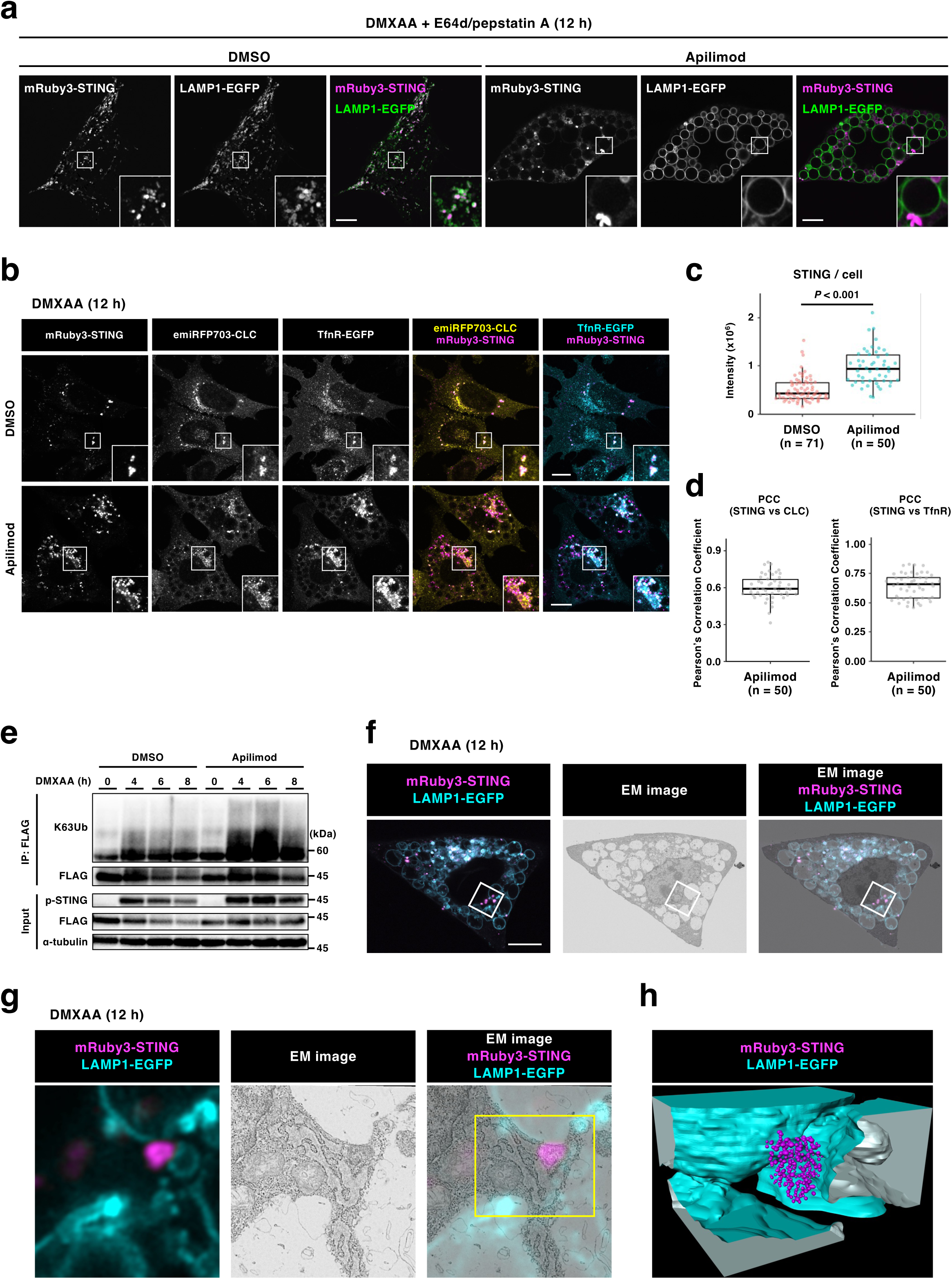
PIKfyve is required for LMA of STING. **a,** Sting^−/−^ MEFs stably expressing mRuby3-STING and LAMP1-EGFP were pretreated with apilimod (1 µM) for 2 h, stimulated with DMXAA for 12 h in the presence of E64d (30 µM) /pepstatin A (40 µM), and then imaged. **b,** Sting^−/−^ MEFs stably expressing mRuby3-STING, emiRFP703-CLC, and TfnR-EGFP were pretreated with apilimod (1 µM) for 2 h, stimulated with DMXAA for 12 h, fixed, and then imaged. **c,** The fluorescence intensity of mRuby3-STING in cells in (**b**) was quantified. **d,** The Pearson’s correlation coefficient in cells in (**b**) is shown. **e,** Sting^−/−^ MEFs reconstituted with FLAG-STING were pretreated with apilimod (1 µM) for 2 h, and then stimulated with DMXAA for the indicated times. FLAG-STING was immunoprecipitated with anti-FLAG antibody. The cell lysates and the immunoprecipitated proteins were analyzed by western blot. **f,g,** Sting^-/-^ MEFs stably expressing mRuby3-STING (magenta) and LAMP1-EGFP (cyan) were pretreated with apilimod (1 µM)for 2 h, and then stimulated with DMXAA for 12 h. LAMP1-positive lysosomes and STING-positive compartments were identified by Airyscan super-resolution microscopy before processing for scanning EM examination. **h**, 3D rendering of segmented lysosomes (cyan) and STING-positive compartments (magenta) in the area indicated by the yellow box in (**g**). Scale bars, 10 µm.

We have previously shown that STING- and clathrin-positive vesicles of an RE origin are directly encapsulated by lysosomes^13^. To examine whether STING reaches REs and is loaded to clathrin-positive vesicles in apilimod-treated cells, we expressed three fluorescence proteins in Sting^-/-^ MEFs: mRuby3-STING, an RE protein transferrin receptor (TfnR) tagged with EGFP, and emiRFP703-Clathrin light chain (CLC). In control cells, the signal of mRuby3-STING mostly diminished after 12-h stimulation (Fig. 3b). In contrast, the signal of mRuby3-STING was largely restored (Fig. 3b-c), and colocalized with TfnR-EGFP or emiRFP703-CLC in apilimod-treated cells (Fig. 3b, and d). The accumulation of STING at REs/clathrin-positive compartments was also confirmed in Pikfyve-knockdown cells (Fig. S3a-b). STING undergoes K63-ubiquitination at Lys288 at the post-Golgi compartments^13^. By co-immunoprecipitation analysis, we found that STING underwent K63-ubiquitination in apilimod-treated cells, further supporting that STING reached the post-Golgi compartments (Fig. 3e).

We then exploited three-dimensional correlative light and electron microscopy (3D-CLEM) to examine the morphology of the accumulating STING-positive compartments under PIKfyve inhibition. The analysis revealed that the chunky STING-positive compartment next to lysosomes was a cluster of vesicles (Fig. 3f-h). These results suggested that PIKfyve did not interfere with the post-Golgi traffic of STING to the REs/clathrin-positive vesicles and that the impaired STING degradation was due to the defect of LMA.

### CHMP4B/C, the ESCRT-III components essential for STING degradation, localize to lysosomes in a PI(3,5)P_2_**-**dependent manner

ESCRT is the evolutionarily conserved multimeric protein complex that deforms various organelle membranes. Given that LMA degradation of STING requires the function of ESCRT^13, 22, 23^, we hypothesized that some subunits of the ESCRT localize at lysosomes and act as a PI(3,5)P_2_ effector. To test this hypothesis, we screened all the mammalian ESCRT and the associated proteins for STING degradation with the dual Luciferase reporter system (Fig. 1a). We found that knockdown of 10 genes (Tsg101, Vps36, Chmp2a/b, Chmp4b/c, Ist1, Vps4a/b, and Lip5) significantly suppressed the STING degradation, compared to knockdown with siRNA control (Fig. 4a and Fig. S4).

**Fig. 4.**
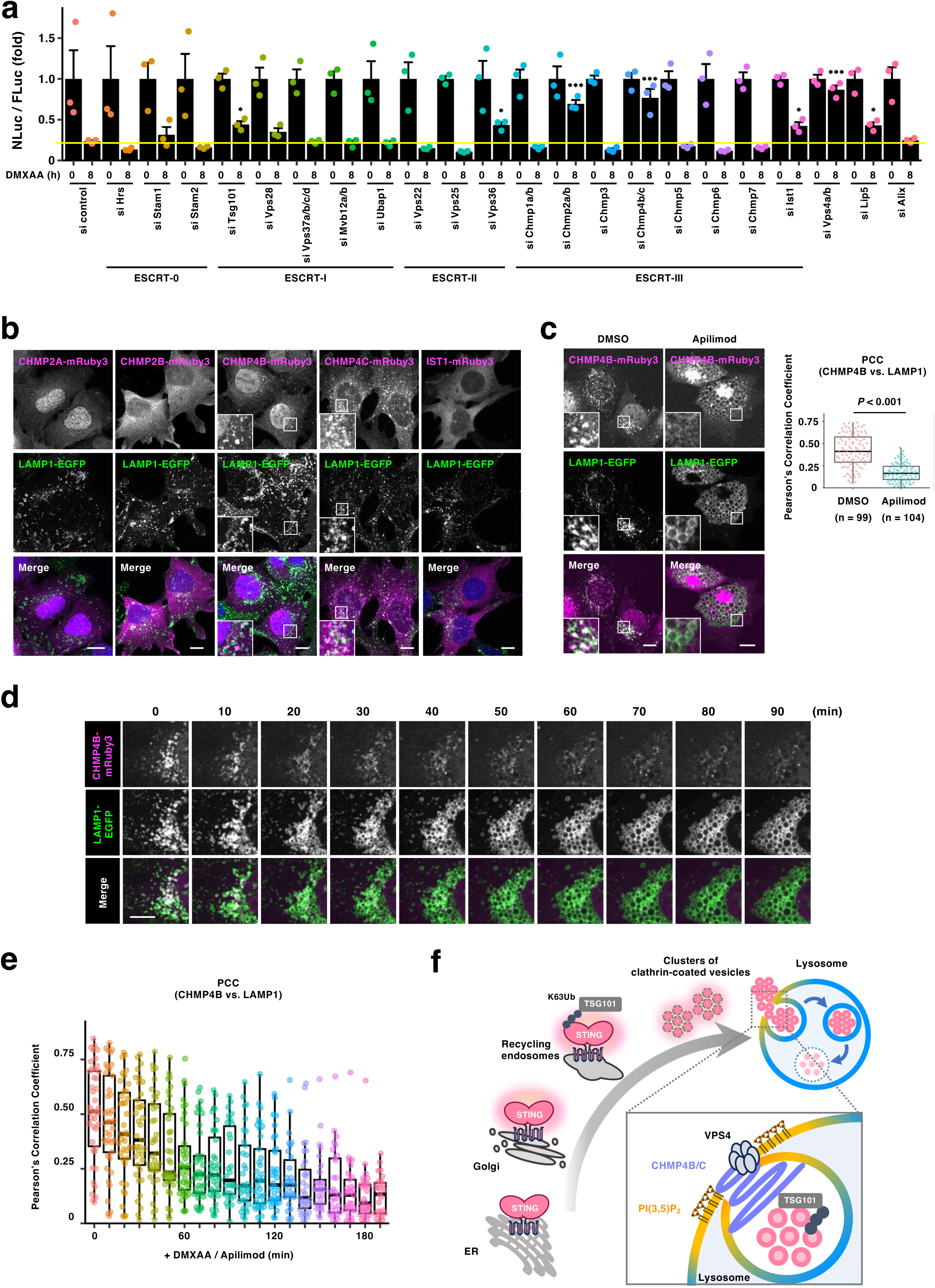
CHMP4B/C localize to lysosomes in a PI(3,5)P_2_-dependent manner. **a,** Sting^-/-^ MEFs stably expressing FLuc-P2A-NLuc-mSTING were treated with the indicated siRNAs, and stimulated with DMXAA for 8 h. The luminescence of FLuc and NLuc was then quantitated, and the NLuc/FLuc ratio was plotted. **b,** DMXAA-stimulated MEFs stably expressing LAMP1-EGFP and the indicated ESCRT-III proteins tagged with mRuby3 were fixed and imaged. **c,** DMXAA-stimulated MEFs stably expressing CHMP4B-mRuby3 and LAMP1-EGFP were treated with apilimod (1 µM) for 3 h, and then imaged. The Pearson’s correlation coefficient between CHMP4B-mRuby3 and LAMP1-EGFP is shown. **d,** MEFs stably expressing CHMP4B-mRuby3 and LAMP1-EGFP were imaged every 10 min after treatment with DMXAA and apilimod (1 µM). **e,** The Pearson’s correlation coefficient between CHMP4B-mRuby3 and LAMP1-EGFP is shown. **f,** A graphical abstract illustrating the function of PI(3,5)P_2_ and CHMP4B/C in LMA. Data are presented as mean ± standard error of the mean. Scale bars, 10 µm.

ESCRT-III forms a neck-like structure on the target membranes, facilitating membrane scission reactions^15, 16^. We thus focused on 5 ESCRT-III genes (Chmp2a/b, Chmp4b/c, and Ist1) out of the 10 genes, and examined their lysosomal localization. As shown (Fig. 4b), mRuby3-tagged CHMP4B and CHMP4C co-localized at LAMP1-EGFP, whereas mRuby3-tagged CHMP2A, CHMP2B, and IST1 localized diffusively throughout the cytoplasm. These results suggested a role of CHMP4B/CHMP4C in LMA. Of note, knockdown of Chmp4b and Chmp4c resulted in the accumulation of mRuby3-STING colocalized with TfnR and clathrin (Fig. S5).

Intriguingly, we found that the lysosomal localization of CHMP4B was lost with apilimod treatment (Fig. 4c). This finding was further corroborated by the super-resolution live cell imaging (Fig. 4d-e and Fig. S6). The lysosomal localization of CHMP4C was also lost with apilimod treatment (Fig. S7). These results suggested that CHMP4B/CHMP4C serve as a lysosomal PI(3,5)P_2_ effector, which facilitates LMA degradation of STING (Fig. 4f).

## Discussion

There are five ways for the degradative substrates to get into the interior space or lumen of lysosomes in mammals^14^. These are namely: endocytic pathway, macroautophagy, chaperon-mediated autophagy, endosomal microautophagy (EMA), and LMA. Among them, EMA and LMA are cognate processes, both exploiting the ESCRT machinery to encapsulate the degradative substrates with the limiting membrane of endosomes or lysosomes and to generate ILVs. We have previously shown that LMA, not EMA, was required for STING degradation^13^. However, why STING degradation depends exclusively on LMA has not been addressed.

In the present study, we found that a specific set of ESCRT genes was required for STING degradation: ESCRT-0 had a marginal contribution. Five ESCRT-III genes (Chmp2a/b, Chmp4b/c, and Ist1) were essential. In EMA, the recognition of ubiquitinated transmembrane proteins by HRS (a subunit of ESCRT-0) is critical to evoke the sequential recruitment of the other ESCRT subcomplexes. HRS localizes on early endosomes (EEs) through its affinity to PI(3)P, a phosphoinositide enriched in EEs^24, 25^. In this regard, HRS may preferentially bind ubiquitin chain that is placed proximal to EE membrane. In contrast, TSG101 is largely cytosolic^13^. As shown in the previous study, TSG101 was recruited from the cytosol to RE-derived vesicles that harbor K63-ubiquitinated STING, indicating that TSG101 can bind ubiquitin chain at any localization in the cytosol. Therefore, the distinct subcellular localization of HRS and TSG101 may explain, at least in part, the selective engagement of LMA for STING degradation.

The size of the endosomal ILVs is rather small, ranging between 40 and 80 nm^26^, which highly contrasts with that of lysosomal ILVs ranging between 200 and 300 nm^13^. In reconstitution experiments, yeast ESCRT-III proteins cooperate to form neck structure on liposomes. Intriguingly, the inclusion of SNF7 (CHMP4 in mammals) induces an extension of the neck structures into elongated tubes^27^. These findings suggest that specific combinations of ESCRT-III proteins can generate distinct types of neck structure. CHMP2A/B, CHMP4A/B, and IST1, the five components that are engaged in STING degradation, may cooperate to form a neck structure in lysosomal membrane, distinct from the one formed in EE membrane.

Pikfyve knockdown formed dilated lysosomes and endosomes. As LMA exploits the limiting membrane of lysosomes to engulf cytosolic degradative substrates, lysosomal membrane itself is also internalized into lysosomal lumen. In this regard, LMA functions to reduce the surface area of lysosomes. Therefore, impaired LMA may also contribute to the dilation process of lysosomes, as well as the membrane traffic defect^28^. A vacant lumen of lysosomes in apilimod-treated cells endorsed this possibility.

In mice, PIKfyve knockout leads to early embryonic lethality^29^, and tissue-specific knockouts reveal critical roles in myeloid, oligodendrocytes, muscle, and intestine cells ^30, 31^. PIKfyve is thought to exist as a large multi-protein complex with two accessory proteins, VAC14^32^ and FIG4^33^. Recent studies have disclosed involvement of the Pikfyve/Vac14/Fig4 in human diseases^31, 34^. Loss-of-function Fig4 mutations are involved in the amyotrophic lateral sclerosis (ALS), primary lateral sclerosis, and Charcot-Marie-Tooth disorder. Loss-of-function mutations in Vac14 lead to a progressive early-onset neurological disease. Loss-of-function mutation in PIKfyve causes congenital cataracts^35-8^. Given that extensive engagement of STING with inflammatory and neurogenerative diseases including ALS, the prolonged STING signalling may contribute to pathogenesis of these Pikfyve-related diseases.

## Material and method

### Ethical approval

MEFs from mice were collected according to ethics number PA17-84 approved by the Institute of Medical Sciences of the University of Tokyo.

### Reagents

Antibodies used in this study are shown in Supplementary Table 2. The following reagents were purchased from the manufacturers as noted: DMXAA (14617, Cayman), bafilomycin A1 (11038, Cayman), brefeldin A (11861, Cayman), Custom Kinase Screening Library (9003376, Cayman), apilimod (HY-14644, MCH MedChemExpress Co.), Vps34-IN1(17392, Cayman), E64d (4321, Peptide Institute), pepstatin A (4397, Peptide Institute), anti-FLAG M2 Affinity Gel (A2220, Sigma), and HT-DNA (D6898, Sigma).

### Cell culture

MEFs were obtained from embryos of WT or Sting^-/-^ mice at E13.5 and immortalized with SV40 Large T antigen. MEFs were cultured in DMEM supplemented with 10% fetal bovine serum (FBS) and penicillin/streptomycin/glutamine (PSG) in a 5% CO_2_ incubator. MEFs that stably express tagged proteins were established using retrovirus. Plat-E cells were transfected with pMXs vectors, and the medium that contains the retrovirus was collected. MEFs were incubated with the medium and then selected with puromycin (2 µg ml^−1^), blasticidin (4 µg ml^−1^), or hygromycinB (100 µg ml^−1^) for several days. RAW-Lucia ISG-KO-STING Cells (InvivoGen) were cultured in DMEM supplemented with 10% FBS, normocin (100 µg ml^−1^), and PSG.

### Immunocytochemistry

Cells were seeded on coverslips (13 mm No.1 S, MATSUNAMI), fixed with 4% paraformaldehyde (PFA) in PBS at room temperature for 15 min, and permeabilized with digitonin (50 µg ml^−1^) in PBS or 0.1% TritonX-100 in PBS at room temperature for 5 min. After blocking with 3% BSA in PBS, cells were incubated with primary antibodies followed by secondary antibodies at room temperature for 45 min. When necessary, cells were stained with DAPI. Cells were then mounted with ProLong Glass Antifade Mountant (P36984, Thermo Fisher Scientific).

### Confocal microscopy

Confocal microscopy was performed using LSM880 with Airyscan (Zeiss) with 20 × 0.8 Plan-Apochromat dry lens, 63 × 1.4 Plan-Apochromat oil immersion lens or 100 × 1.46 alpha-Plan-Apochromat oil immersion lens. Images were analyzed and processed with Zeiss ZEN 2.3 SP1 FP3 (black, 64 bit) (version 14.0.21.201) and Fiji (version 2.9.0/1.54h). Pearson’s correlation coefficient was quantified by BIOP JACoP in Fiji plugin.

### Nano-Glo^®^ Dual-Luciferase^®^ Reporter assay

Cells in culture medium were lysed using ONE-Glo^TM^ EX Reagent, in a volume equal to the culture medium, at room temperature. After the measurement of the luminescence of FLuc, an equal volume of NanoDLR^TM^ Stop & Glo^®^ Reagent was added to quench the FLuc signal, and the luminescence of NLuc was measured with GloMax Navigator Microplate Luminometer (Promega, version 3.1.0). The luminescence value of NLuc was divided by that of FLuc to obtain the NLuc/FLuc ratio. In Fig. 1f, the NLuc/FLuc ratios obtained from cells treated with inhibitors were normalized to the averaged NLuc/FLuc ratio obtained from cells treated with DMSO. In Fig. 1a, 1e, 1h, 4a, S2c, S2d, and S2e, the NLuc/FLuc ratios obtained from cells stimulated with DMXAA and the corresponding inhibitor or siRNA were normalized to the averaged NLuc/FLuc ratio obtained from unstimulated cells treated with the corresponding inhibitor or siRNA.

### PCR cloning

Complementary DNAs (cDNAs) encoding mouse Sting, mouse Chmp2a, mouse Chmp2b, mouse Chmp4b, mouse Chmp4c, and mouse Ist1 were amplified by PCR with mouse cDNA library. The amplified products were inserted into pMX-IPuro-NLuc-MCS-P2A-FLuc, pMX-IPuro-MCS-NLuc-P2A-FLuc, pMX-IPuro-FLuc-P2A-NLuc-MCS, pMX-IPuro-FLuc-P2A-MCS-NLuc, or pMXs-IBla-MCS-mRuby3. The plasmids encoding NLuc and FLuc were purchased from Promega.

### Type I interferon bioassay

Cell culture supernatants of MEFs stimulated with DMXAA were added to Raw264.7-Lucia ISG-KO-STING Cells (Invivogen). Ten hours after incubation, the luciferase activity was measured by GloMax Navigator Microplate Luminometer (Promega, version 3.1.0).

### qRT-PCR

Total RNA was extracted from cells using SuperPrep II (TOYOBO) or ISOGEN-II (Nippon gene), and reverse-transcribed using ReverTraAce qPCR RT Master Mix with gDNA Remover (TOYOBO). Quantitative real-time PCR (qRT-PCR) was performed using KOD SYBR qPCR (TOYOBO) and LightCycler 96 (Roche). Target gene expression was normalized to the expression of Gapdh.

### Western blotting

Proteins were separated in polyacrylamide gel and then transferred to polyvinylidene difluoride membranes (Millipore). These membranes were incubated with primary antibodies, followed by secondary antibodies conjugated to peroxidase. The proteins were visualized by enhanced chemiluminescence using Fusion SOLO.7S.EDGE (Vilber-Lourmat).

### Immunoprecipitation

Cells were washed with ice-cold PBS and scraped in immunoprecipitation buffer composed of 50 mM HEPES-NaOH (pH 7.2), 150 mM NaCl, 5 mM EDTA, 1% Triton X-100, protease inhibitor cocktail (Nacalai Tesque, 25955, dilution 1:100) and phosphatase inhibitors (8 mM NaF, 12 mM β-glycerophosphate, 1 mM Na_3_VO_4_, 1.2 mM Na_2_MoO_4_, 5 mM cantharidin, and 2 mM imidazole). The cell lysates were centrifuged at 15,000 rpm for 10 min at 4 ℃, and the resultant supernatants were pre-cleared with Ig-Accept Protein G (Nacalai Tesque) at 4 °C for 30 min. The lysates were then incubated overnight at 4 ℃ with anti-FLAG M2 Affinity Gel. The beads were washed four times with immunoprecipitation wash buffer (50 mM HEPES-NaOH (pH 7.2), 150 mM NaCl, and 0.1% Triton X-100) and eluted with the Laemmli sample buffer.

### RNA interference

siRNAs used in the present study were purchased from Dharmacon or Thermo Fisher Scientific. Cells were transfected with siRNAs (5 nM or 20 nM) using Lipofectamine RNAiMAX (Invitrogen) according to the manufacturer’s instruction.

### 3D-CLEM

Cells on coverslips were fixed with 2% glutaraldehyde in 0.1 M phosphate buffer (pH 7.4) for 24 h at 4 ℃, post-fixed with 1% osmium tetroxide (0.1 M PB) for 1 h at 4 ℃, and rinsed with Milli-Q water. The cells were then stained with a 1% aqueous solution of uranyl acetate for 1 h, dehydrated through a graded series of ethanol, embedded in Epoxy resin, and incubated for 48 h at 60 ℃. After resin polymerization, the coverslip was removed from the resin on a hot plate for 1 min at 100 ℃. Cells previously imaged by light microscopy were identified by their position on the grid. A 0.5 mm^2^-area containing cells of interest was trimmed with a razor blade. 150 consecutive ultrathin sections (60 nm thickness) were cut with an ultramicrotome (EM UC7, Leica) using a diamond knife (Diatome), mounted onto a glass slide, stained with a 1% aqueous solution of uranyl acetate for 10 min and with a 1% aqueous solution of lead citrate for 5 min. Backscattered electron images of these serial sections were obtained with a scanning electron microscope (SU-70, Hitachi). CLEM images were constructed using image processing software (Photoshop CC, Adobe Systems Inc.). 3D rendering of the region of interest was performed with Amira 3D (Thermo Fisher Scientific)^39^.

### RNA-sequencing analysis

Total RNA was extracted by primary MEFs. RNA sequencing was performed using Novaseq 6000 (Illumina, Inc., CA, USA) by the NGS core facility at the Research Institute for Microbial Diseases of Osaka University. Estimation of expression counts was executed by kallisto (ver. 0.50.1)^40^ using “Mus_musculus.GRcm38.cdna.all.fa”. The expression counts were normalized as scaled transcript per million (TPM) by tximport package (ver. 1.26.1) in R (ver. 4.3.2). Signature scores of inflammation-related gene sets were calculated in GSVA (ver. 1.44.5) using the “ssGSEA” method^41^. Heatmap was drawn using Morpheus (Broad Institute, https://software.broadinstitute.org/morpheus/).

### Statistics and reproducibility

Error bars displayed in bar plots throughout the present study represent the standard error of the mean (SEM) unless otherwise indicated. The SEM was calculated from at least triplicate samples. In box-and-whisker plots, the box bounds the interquartile range divided by the median, one-way analysis of variance followed by Tukey-Kramer post hoc test for multiple comparisons (Figs. 1e, 1h, 2b, 2d, 2f, 3c, 4c, S2c-e, and S4), or Dunnett’s test for multiple comparisons (Fig. 4a) with R (version 4.1.2) and KNIME (version 4.5.1); *P < 0.05; ***P < 0.001; NS not significant (P > 0.05). No statistical method was used to pre-determine sample size. No data were excluded from the analyses. The experiments were not randomized.

---

**Fig. S1.**
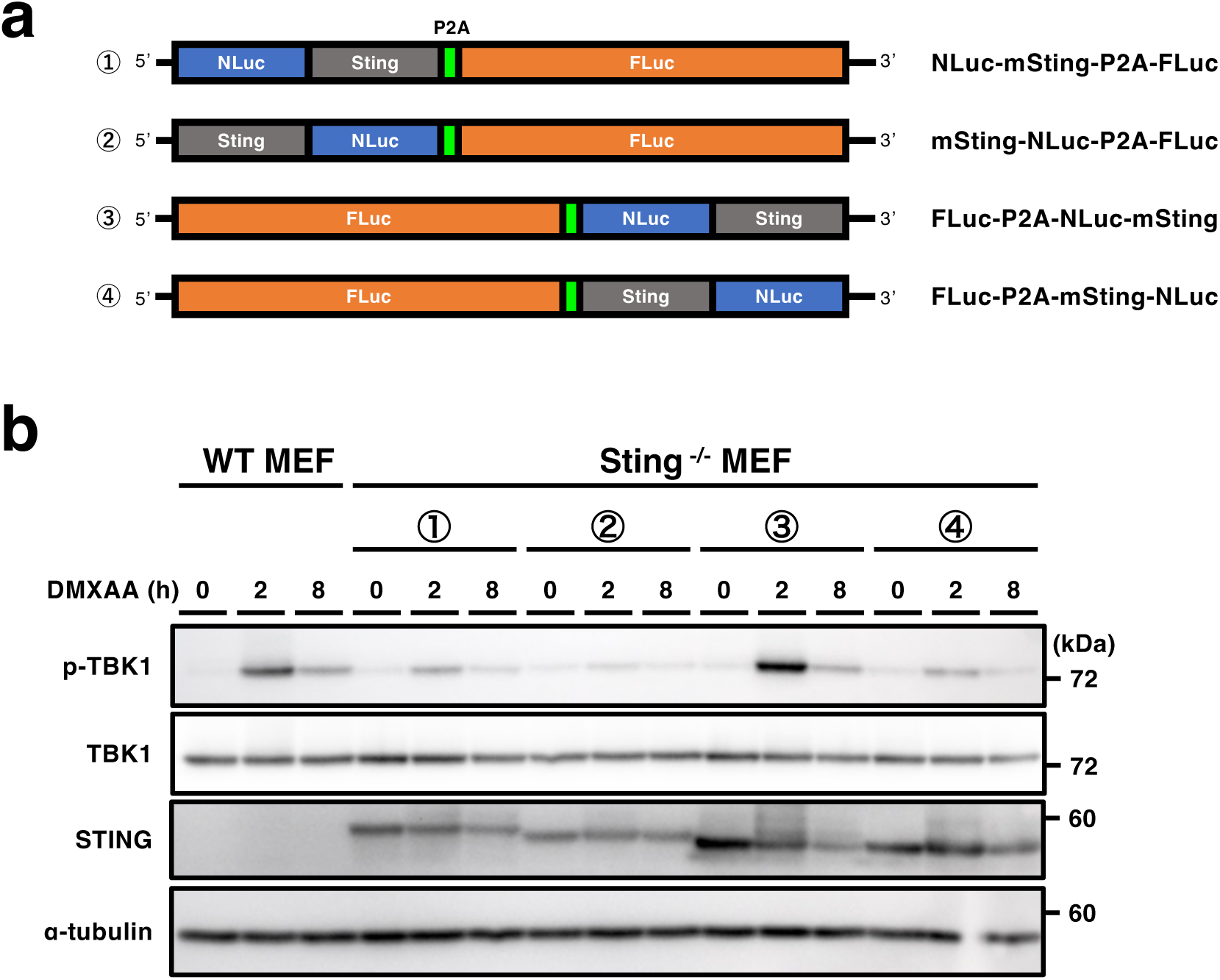
**a**, Schematic illustration of the plasmids encoding NLuc-mSting-P2A-FLuc, mSting-NLuc-P2A-FLuc, FLuc-P2A-NLuc-mSting, and FLuc-P2A-mSting-NLuc. **b**, Cells expressing the individual construct shown in (**a**) were stimulated DMXAA for the indicated times. Cell lysates were analyzed by western blot.

**Fig. S2.**
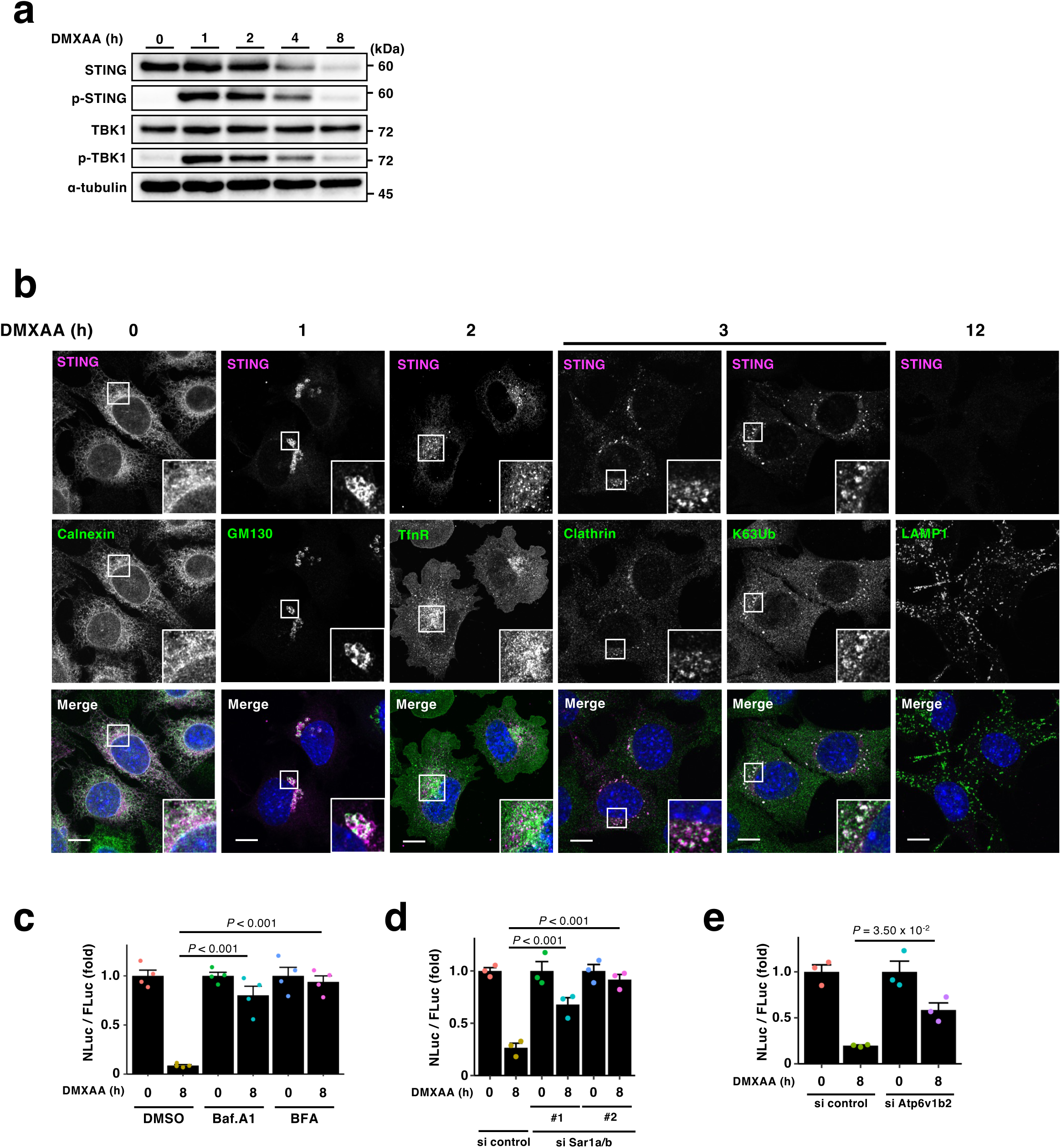
**a,** FLuc-P2A-NLuc-mSTING was stably expressed in Sting^-/-^ MEFs. Cells were stimulated with DMXAA for the indicated times. Cell lysates were analyzed by western blot. **b,** Sting^-/-^ MEFs stably expressing FLuc-P2A-NLuc-mSTING were treated with DMXAA for the indicated times. Cells were immunostained with anti-LgBiT, anti-STING, anti-Calnexin, anti-GM130, anti-TfnR, anti-Clathrin, anti-K63 Ubiquitin, or anti-Lamp1 antibody. **c,** Sting^-/-^ MEFs stably expressing FLuc-P2A-NLuc-mSTING were stimulated with DMXAA for 8 h in the presence of Baf.A1 (100 nM) or BFA (3 µg/ml). The luminescence of FLuc and NLuc was quantitated, and the NLuc/FLuc ratio was plotted. **d, e,** Sting^-/-^ MEFs stably expressing FLuc-P2A-NLuc-mSTING were treated with the indicated siRNAs for 64 h and then stimulated with DMXAA for 8 h. The luminescence of FLuc and NLuc was quantitated, and the NLuc/FLuc ratio was plotted. Data are presented as mean ± standard error of the mean. Scale bars, 10 µm.

**Fig. S3.**
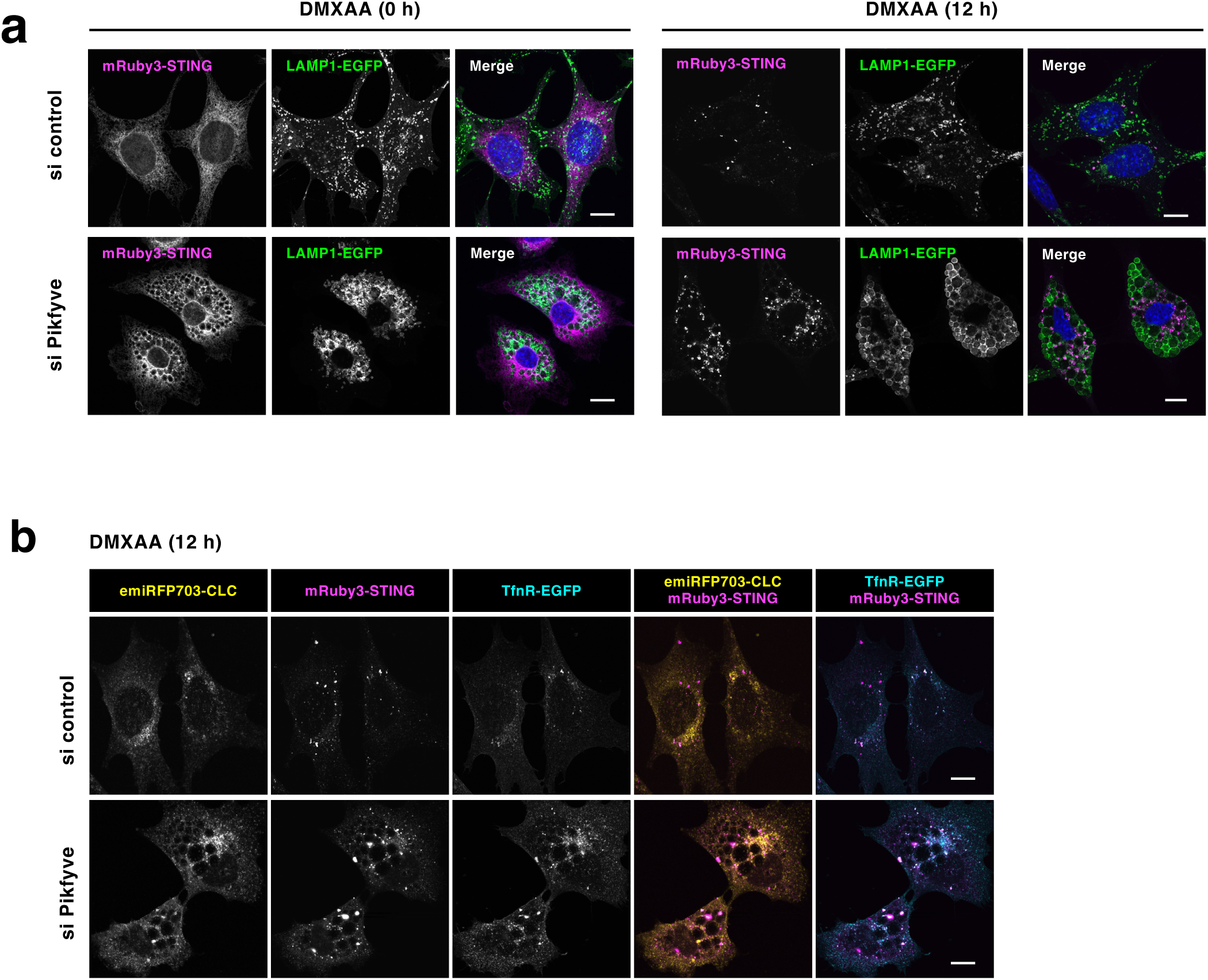
**a,** Sting^-/-^ MEFs stably expressing mRuby3-mSTING and LAMP1-EGFP were treated with the indicated siRNAs and then stimulated with DMXAA for 12 h. **b,** Sting^-/-^ MEFs stably expressing mRuby3-mSTING, emiRFP703-CLC, and TfnR-EGFP were treated with the indicated siRNAs and then stimulated with DMXAA for 12 h. Scale bars, 10 µm.

**Fig. S4.**
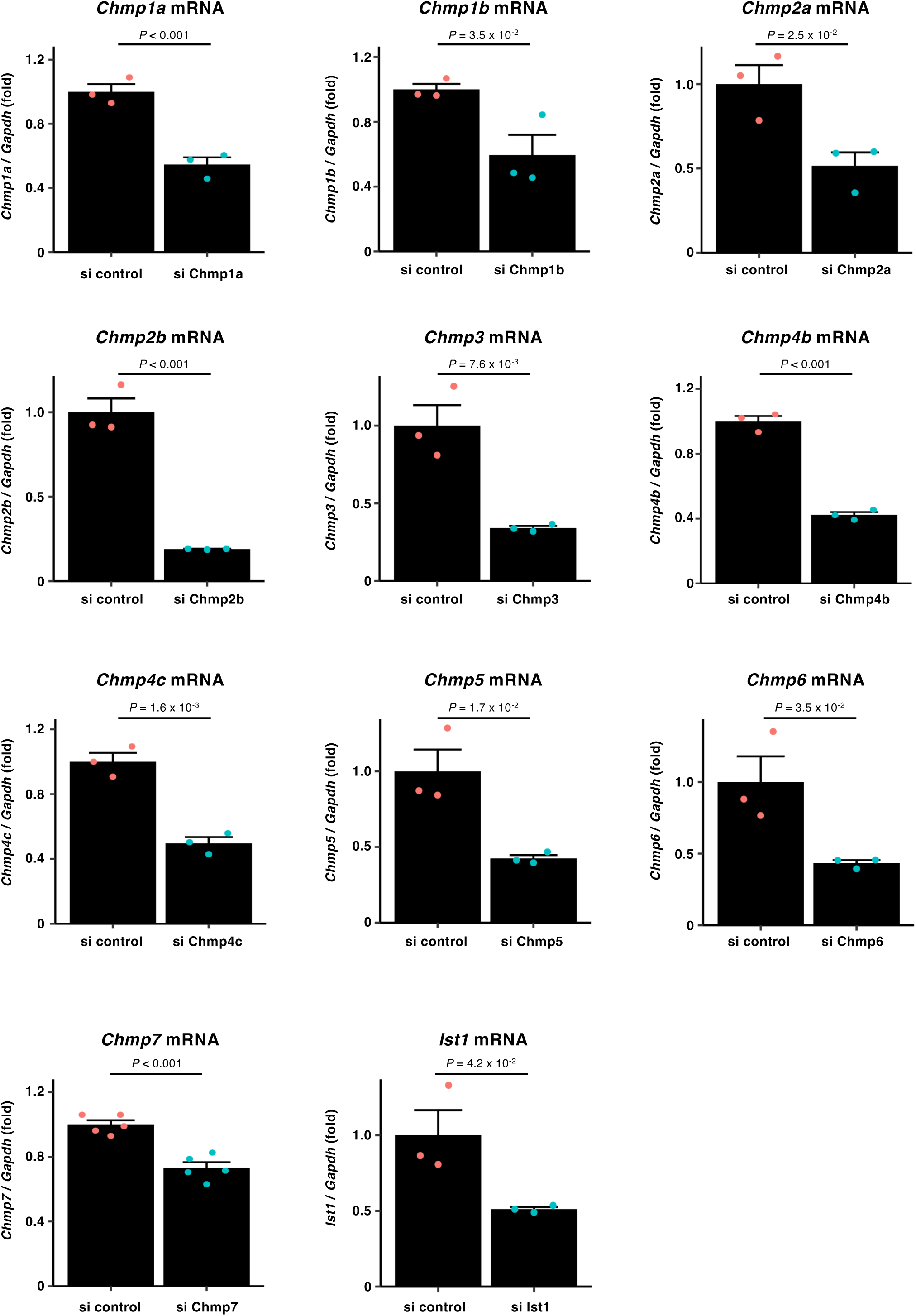
Cells were treated with the indicated siRNAs for 72 h. The knockdown efficiency of the individual genes was quantitated with qRT-PCR. Data are presented as mean ± standard error of the mean.

**Fig. S5.**
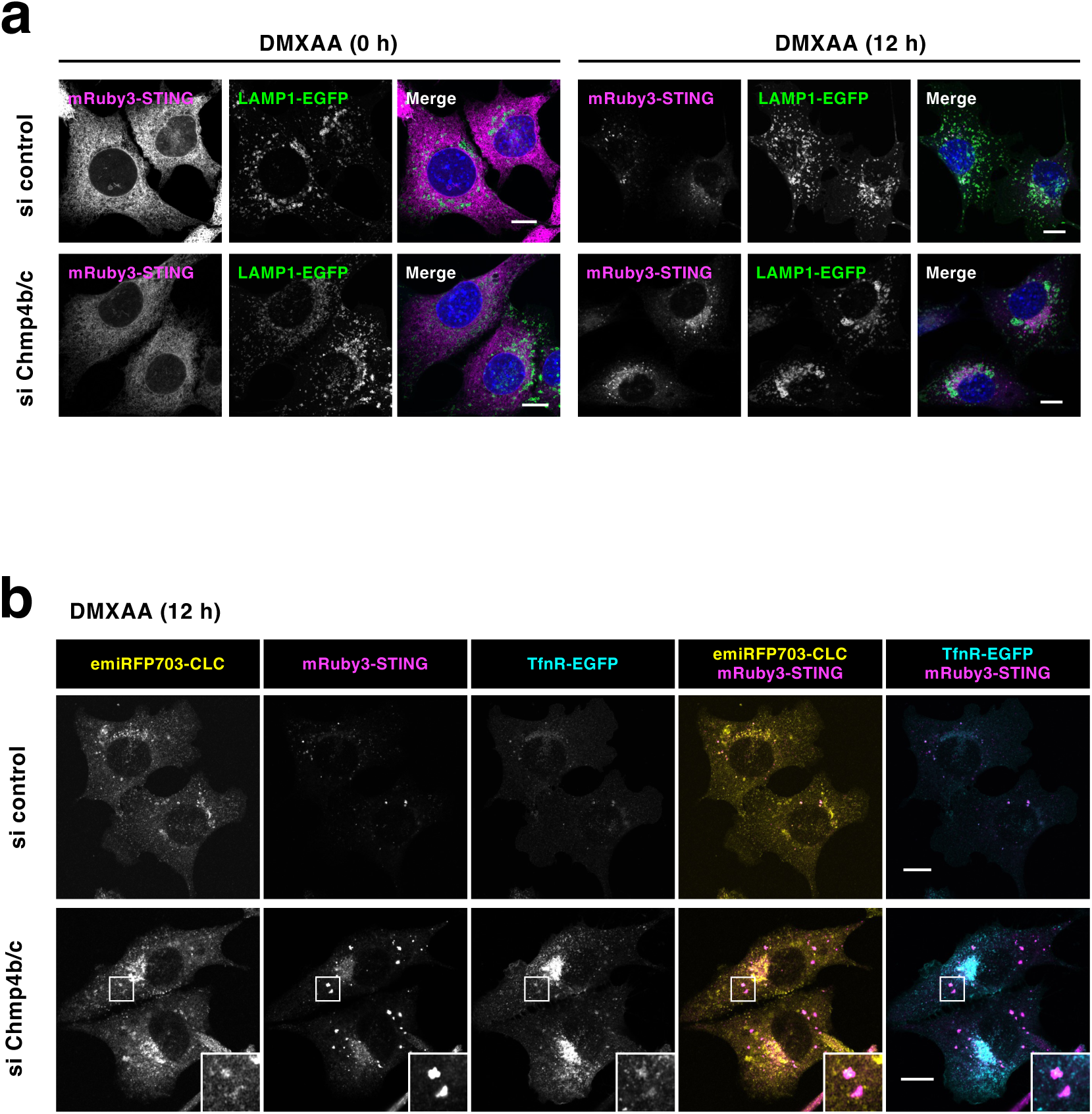
**a,** Sting^-/-^ MEFs stably expressing mRuby3-mSTING and LAMP1-EGFP were treated with the indicated siRNAs and then stimulated with DMXAA for 12 h. **b,** Sting^-/-^ MEFs stably expressing mRuby3-mSTING, emiRFP703-CLC, and TfnR-EGFP were treated with the indicated siRNAs and then stimulated with DMXAA for 12 h. Scale bars, 10 µm.

**Fig. S6.**
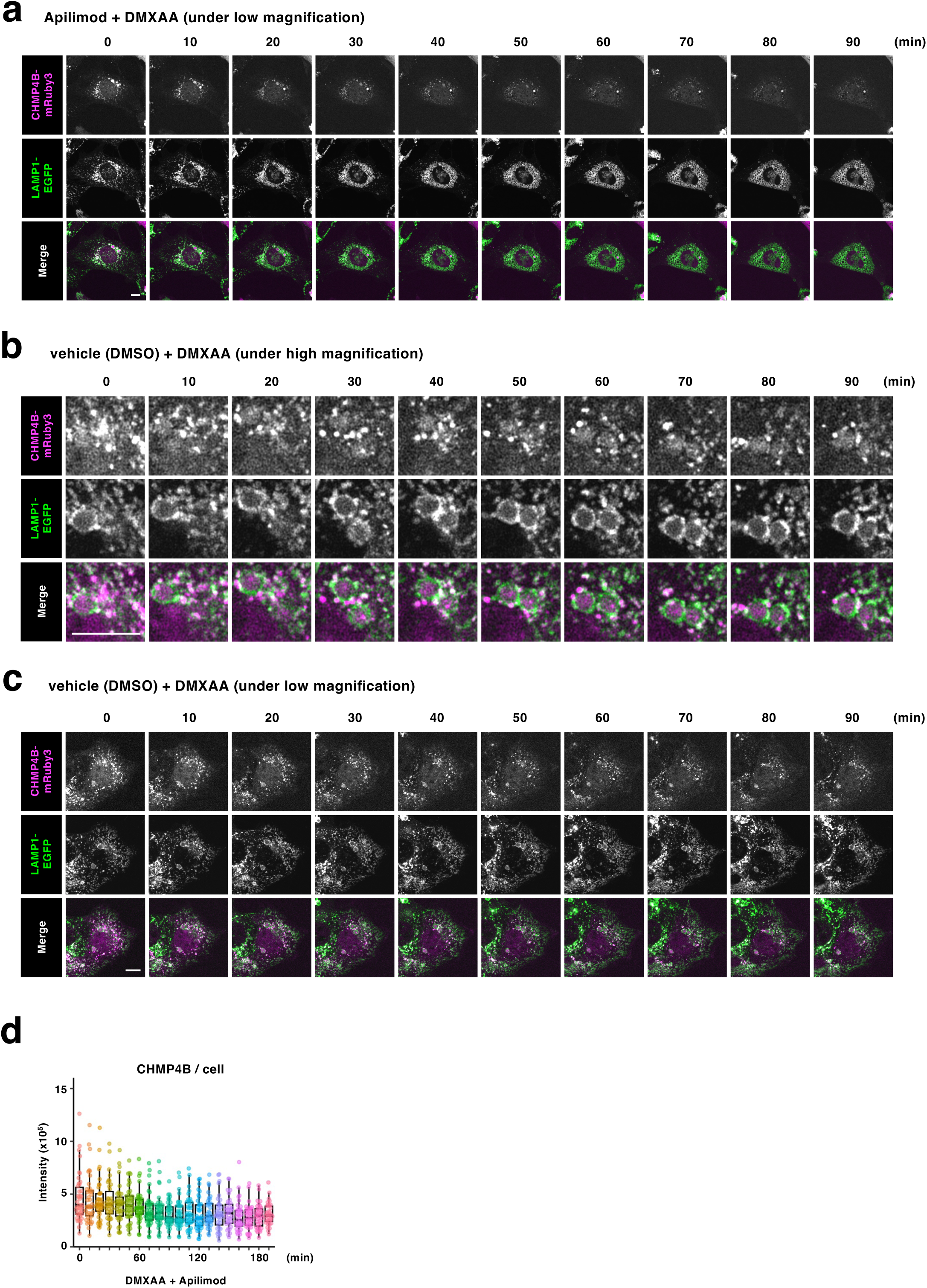
**a-c,** MEFs stably expressing CHMP4B-mRuby3 and LAMP1-EGFP were imaged every 10 min after treatment with apilimod (1 µM) or DMSO in the presence of DMXAA. **d**, The fluorescence intensity of CHMP4B-mRuby3 in cells in (**a**) was quantified. Scale bars, 10 µm.

**Fig. S7.**
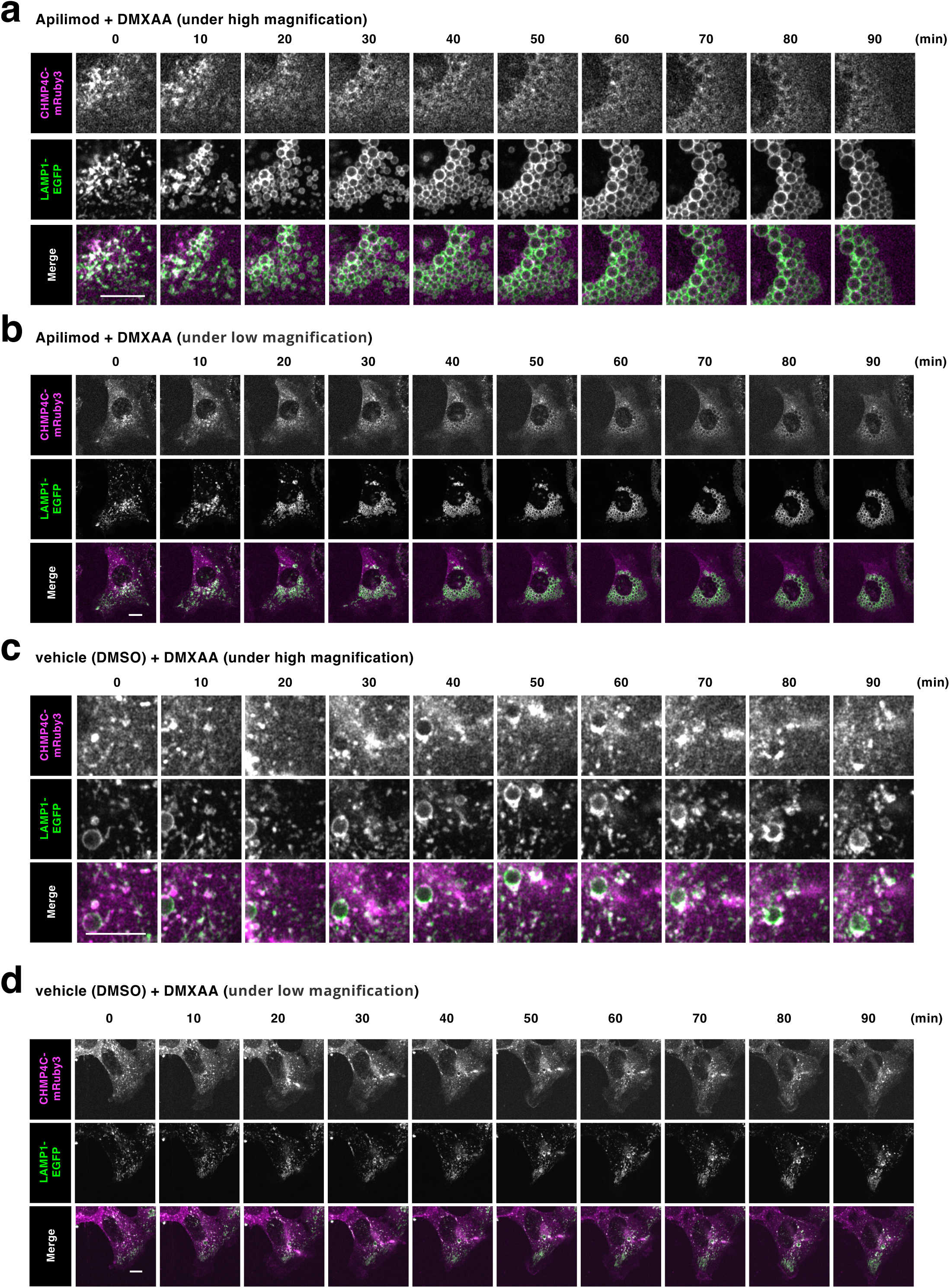
**a-d,** MEFs stably expressing CHMP4C-mRuby3 and LAMP1-EGFP were imaged every 10 min after treatment with apilimod (1 µM) or DMSO in the presence of DMXAA. Scale bars, 10 µm.

| Inhibitor | Target | Nluc/Fluc(fold) |
| --- | --- | --- |
| Staurosporine | PKC | 91.25 |
| YM-201636 | PIKfyve | 42.03 |
| Baf A1 |  | 34.92 |
| SC-1 | ERK1 | 22.19 |
| AZD 7762 | Chk1, Chk2 | 14.93 |
| PIK-75<br>(hydrochloride) | PI3K p110α | 11.38 |
| BIBF 1120 | Multi-kinase | 10.61 |
| PI-103 | DNA-PK, PI3K,<br>mTOR | 6.08 |
| GSK1059615 | PI3Kα | 4.22 |
| URMC-099 | LRRK2, MLKs | 4.09 |
| CAY10626 | PI3Kα, mTOR | 3.71 |
| INK128 | mTOR | 3.68 |
| UNC569 | Axl family | 3.59 |
| R406 | Syk | 3.36 |
| Torin 1 | mTOR | 2.82 |
| PD 173074 | FGFR1 | 2.56 |
| NVP-AEW541<br>(hydrochloride) | IGF-1R | 2.50 |
| TWS119 | GSK3β | 2.49 |
| BI-6727 | PIks | 2.41 |
| JNJ-10198409 | PDGFR | 2.17 |
| CAY10576 | IKKε | 2.07 |
| NVP-TAE226 | FAK, PYK2β | 2.07 |
| HA-1077<br>(hydrochloride) | ROCK- II | 2.02 |
| Dasatinib | Src | 1.93 |
| FRAX597 | PAK1, PAK2,<br>PAK3 | 1.88 |
| RG-13022 | EGFR | 1.77 |
| LFM-A13 | BTK | 1.73 |
| TAS 120 | FGFR family | 1.72 |
| ARQ-092 | PKB/Akt | 1.72 |
| AG-879 | BMX | 1.68 |
| Erbstatin<br>analog | EGFR | 1.61 |
| AS-605240 | PI3Kγ | 1.59 |
| LY2606368 | Chk1 | 1.58 |
| 3-Methyladenine | PI3K | 1.53 |
| AZD 0156 | ATM | 1.50 |
| Ruxolitinib | JAK1, JAK2 | 1.50 |
| 5-Iodotubercidin | CK1, ERK2, PKC | 1.49 |
| BMS-536924 | IGF-1R, InsR | 1.49 |
| AG-1478 | EGFR | 1.46 |
| WHI-P131 | JAK3 | 1.40 |
| JNK Inhibitor XVI | JNK1, JNK2, JNK3, JNKs | 1.39 |
| AG-370 | PDGFR | 1.38 |
| (R)-Crizotinib | c-Met, ALK5 | 1.35 |
| NH125 | EEF2K | 1.28 |
| BMS 345541 (trifluoroacetate salt) | IKKα, IKKβ | 1.28 |
| PHA-665752 | c-Met | 1.26 |
| ZM 447439 | Aurora B | 1.24 |
| AG-825 | ErbB2 | 1.22 |
| CAY10621 | SPHK1 | 1.21 |
| BMS-777607 | Axl family, Met family | 1.19 |
| Canertinib (hydrochloride) | EGFR | 1.18 |
| BX-912 | PDK1 | 1.15 |
| AS-703026 | MEK1, MEK2 | 1.14 |
| LDN-211904 | EphB3 | 1.13 |
| PI3-Kinase $\alpha$ Inhibitor 2 | PI3Kα | 1.12 |
| CAY10505 | CK2 | 1.10 |
| BGJ398 | FGFR family | 1.09 |
| XMD16-5 | TNK2 | 1.09 |
| AZD 1208 | PIMs | 1.07 |
| LCK Inhibitor | LcK | 1.07 |
| PP242 | mTOR | 1.07 |
| Tilfrinib | BRK/PTK6 | 1.05 |
| PD 169316 | p38 MAPK | 1.02 |
| TGX-221 | PI3K p110β | 1.02 |
| PD 0325901 | MEK | 1.01 |
| control |  | 1.00 |
| Wortmannin | PI3K | 0.99 |
| ML-9 | Multi-kinase | 0.98 |
| PF-562271<br>(besylate) | FAK | 0.98 |
| KW 2449 | FLT3, Abl | 0.98 |
| WZ4003 | NUAK1,<br>NUAK2 | 0.96 |
| PF-06463922 | ALK | 0.95 |
| Afuresertib<br>(hydrochloride) | PKB/Akt | 0.95 |
| AG-183 | EGFR | 0.94 |
| GSK2334470 | PDK1 | 0.94 |
| AP26113 | ALK | 0.94 |
| D 4476 | CK1 | 0.94 |
| Imatinib<br>(mesylate) | PDGFR, c-Kit | 0.93 |
| H-89<br>(hydrochloride) | Multi-kinase | 0.93 |
| CGP 57380 | MNK1 | 0.92 |
| Mps1-IN-1 | Mps1 | 0.91 |
| GNE-7915 | LRRK2 | 0.91 |
| JNK Inhibitor<br>XVI | DYRK1B | 0.90 |
| PHA-767491<br>(hydrochloride) | Cdc7 | 0.90 |
| H-8<br>(hydrochloride) | PKA, PKG | 0.89 |
| PD 184161 | MEK1, MEK2 | 0.89 |
| MLCK Inhibitor<br>Peptide 18 | MLCK | 0.89 |
| SB-431542<br>(hydrate) | ALK5 | 0.89 |
| AG-1296 | PDGFR | 0.89 |
| Olomoucine | CDKs | 0.88 |
| SB 202190 | p38 MAPK | 0.88 |
| TG003 | CLK1, CLK4 | 0.88 |
| NU 7026 | DNA-PK | 0.88 |
| Erlotinib | EGFR | 0.87 |
| AG-490 | JAK2 | 0.87 |
| KRN 633 | VEGFR family | 0.87 |
| BI-D1870 | RSKs | 0.87 |
| SB 203580 | p38 MAPK | 0.87 |
| Bisindolylmalei<br>mide I | PKC | 0.87 |
| LOXO-195 | Trk family | 0.87 |
| GNF-5 | Bcr-Abl | 0.87 |
| NU 6102 | DNA-PK | 0.86 |
| H-9<br>(hydrochloride) | PKG | 0.86 |
| Tie2 Kinase<br>Inhibitor | Tie2 | 0.85 |
| Leelamine | PDK | 0.84 |
| Bisindolylmalei<br>mide IX<br>(mesylate) | GSK3, PKC | 0.84 |
| Syk Inhibitor II | Syk | 0.84 |
| KN-62 | CAMK II | 0.83 |
| Apatinib | VEGFR2 | 0.82 |
| KN-93 | CAMK II | 0.82 |
| Sphingosine<br>Kinase<br>Inhibitor 2 | SPHK1 | 0.82 |
| ABT-869 | VEGFR family,<br>PDGFR family | 0.82 |
| CAY10571 | p38α MAPK | 0.82 |
| SC-514 | IKK2 | 0.81 |
| PLX4032 | B-RAF | 0.81 |
| Gö 6983 | PKC | 0.81 |
| (R)-Roscovitine | CDKs | 0.81 |
| AZ 3146 | Mps1 | 0.80 |
| CHIR99021 | GSK3 | 0.80 |
| Nilotinib | Bcr-Abl | 0.80 |
| 17β-hydroxy<br>Wortmannin | PI3K | 0.80 |
| BMS-5 | LIMK1, LIMK2 | 0.79 |
| CAY10561 | ERK2 | 0.78 |
| GW 5074 | Raf-1 | 0.78 |
| SU 6668 | Aurora B | 0.78 |
| Lavendustin C | EGFR | 0.77 |
| Vatalanib<br>(hydrochloride) | VEGFR family | 0.77 |
| LY364947 | ALK5 | 0.76 |
| CAY10622 | ROCK- I ,<br>ROCK- II | 0.76 |
| U-0126 | MEK1, MEK2 | 0.76 |
| Chelerythrine<br>(chloride) | PKC | 0.76 |
| AS-041164 | PI3Kγ | 0.76 |
| Y-27632<br>(hydrochloride) | ROCK- I | 0.75 |
| (S)-H-1152<br>(hydrochloride) | ROCK | 0.75 |
| SP 600125 | JNK1, JNK2,<br>JNK3, JNKs | 0.75 |
| RPI-1 | RET | 0.74 |
| VX-702 | p38 MAPK | 0.74 |
| CAY10578 | CK2 | 0.74 |
| SU6656 | Yes | 0.74 |
| Necrostatin-1 | RIP1 | 0.73 |
| SB-216763 | GSK3 | 0.72 |
| Phthalazinone<br>pyrazole | Aurora A | 0.72 |
| SMI-4a | PIMs | 0.70 |
| Emodin | CK2 | 0.69 |
| LOXO-101 | Trk family | 0.69 |
| Doramapimod | p38 MAPK | 0.67 |
| AG-494 | EGFR | 0.66 |
| Sorafenib | Raf-1 | 0.65 |
| (S)-Glycyl-H-1152<br>(hydrochloride) | ROCK- II | 0.64 |
| Bisindolylmalei<br>mide V | S6K | 0.63 |
| CRT0066101<br>(hydrochloride) | PKD | 0.61 |
| CAY10574 | Cdk9 | 0.50 |
| BIO | GSK3 | 0.39 |
| Kenpaullone | CDKs, GSK3β | 0.35 |
| Indirubin-3'-<br>monoxime | GSK3β | 0.35 |

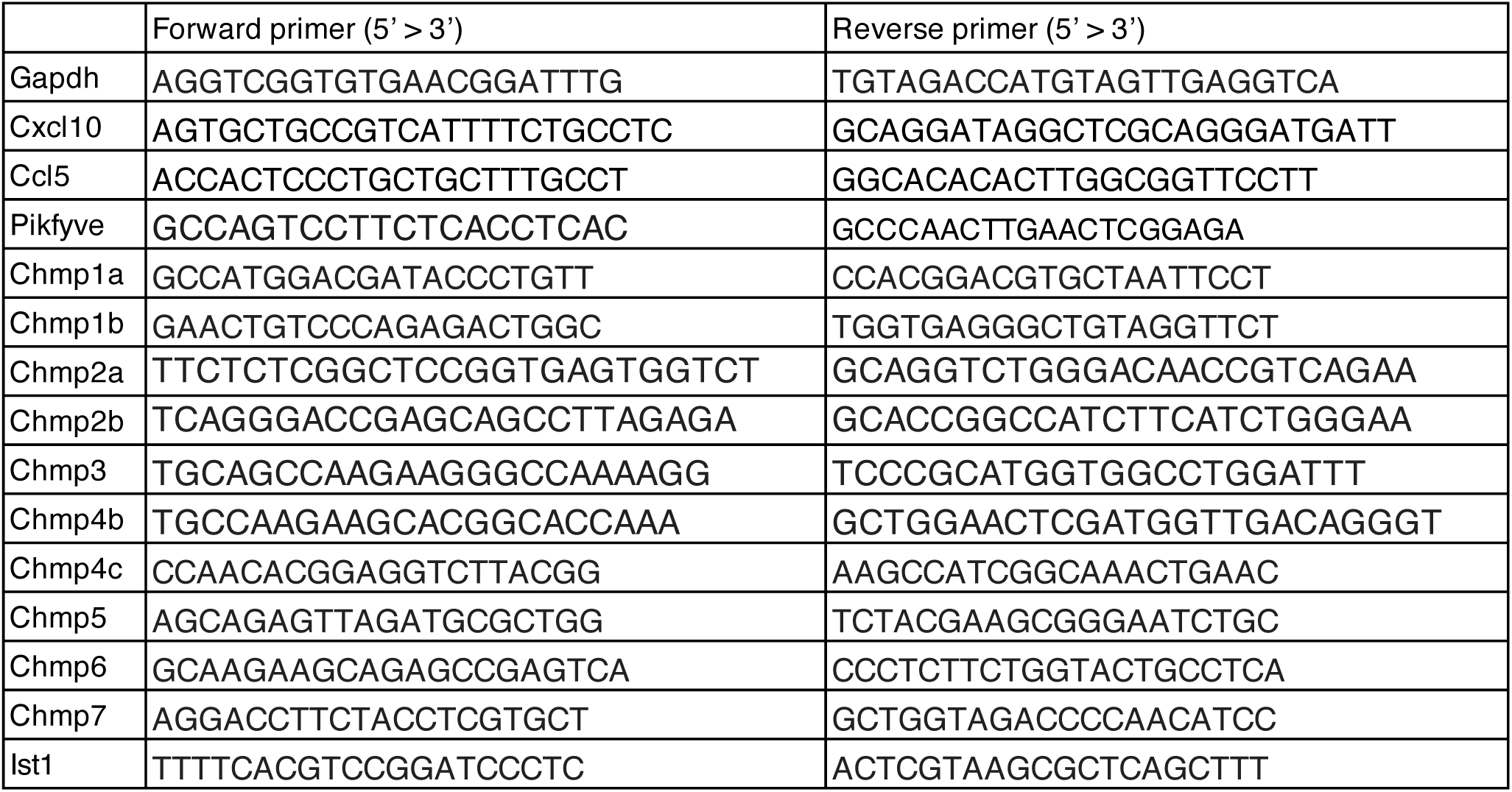

| Protein | Company | Product number | Dilution | Purpose |
| --- | --- | --- | --- | --- |
| STING | proteintech | 19851-1-AP | 1/1000 | WB |
| pSTING | Cell Signaling Technology | D8F4W | 1/1000 | WB |
| α-tubulin | Fujifilm-Wako | 017-25031 | 1/500 | WB |
| GM130 | BD Biosciences | 610823 | 1/1000 | IF |
| pTBK1 | Cell Signaling Technology | D52C2 | 1/1000 | IF/WB |
| TBK1 | Abcam | ab40676 | 1/1000 | WB |
| K63-linked ubiquitin | Merck Millipore | 05-1308 | 1/1000 | IF |
| K63-linked ubiquitin | Abcam | ab179434 | 1/250 | WB |
| Goat Anti-Rabbit IgG (H + L) Mouse/Human ads-HRP | Southern Biotech | 4050-05 | 1/10000 | WB |
| Goat Anti-Mouse IgG (H + L) Human ads-HRP | Southern Biotech | 1031-05 | 1/10000 | WB |
| Donkey Anti-Goat IgG(H+L) Multi-Species SP ads-HRP | Southern Biotech | 6425-05 | 1/10000 | WB |
| Alexa 488-, 568-, or 594- conjugated secondary antibodies | Thermo Fisher Scientific | A21202, A21206, A21207, A21208, A10037 | 1/1000 | IF |
| LC3 | MBL Life science | PM036 | 1/1000 | IF/WB |
| Clathrin heavy chain | Cell Signaling Technology | D3C6 | 1/100 | IF |
| LAMP1 | Invitrogen | 14-1071-82 | 1/1000 | IF |
| CALNEXIN | proteintech | 10427-2-AP | 1/500 | IF |
| PIKfyve | proteintech | 13361-1-AP | 1/1000 | WB |
| FLAG | Bethyl Laboratories | A190-101A | 1/1000 | WB |
| TfnR | Santa Cruz Biotechnology | sc-65882 | 1/100 | IF |
| LgBiT | Promega | N710A | 1/100 | IF |

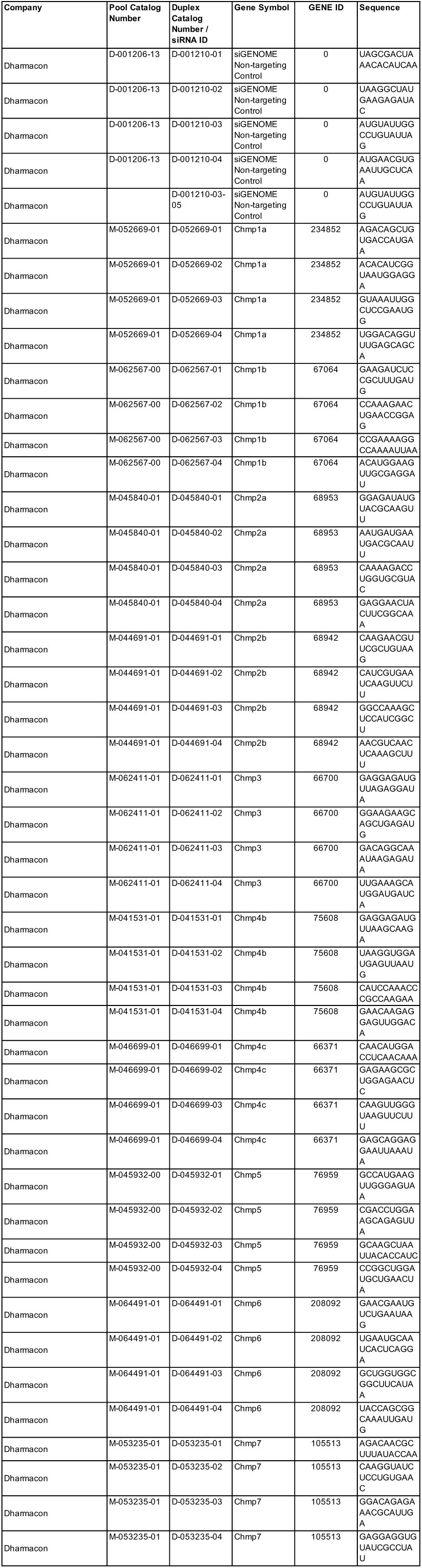
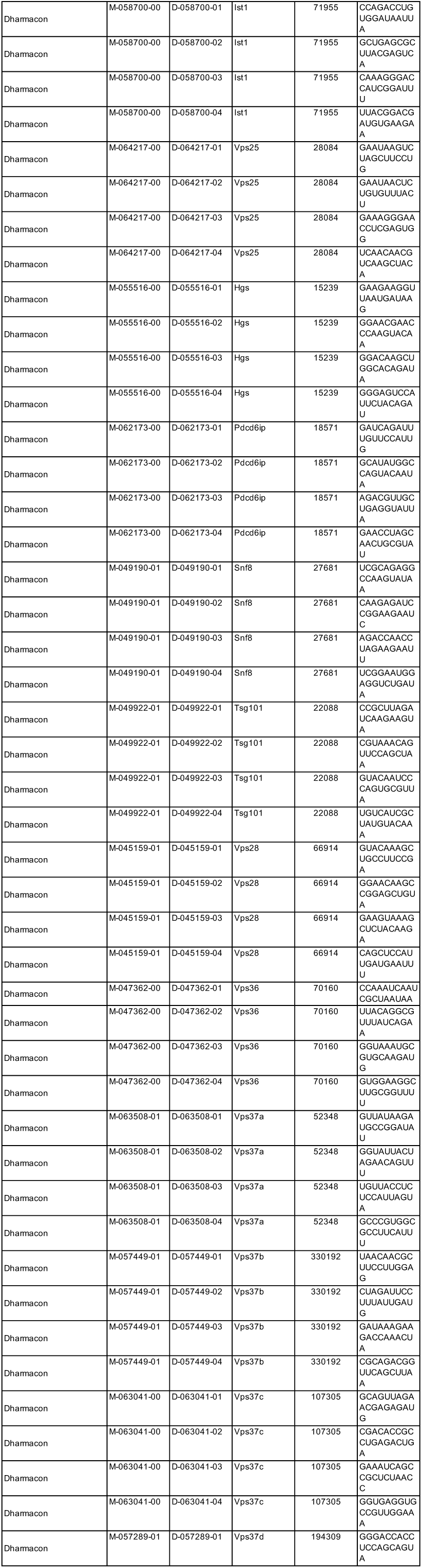
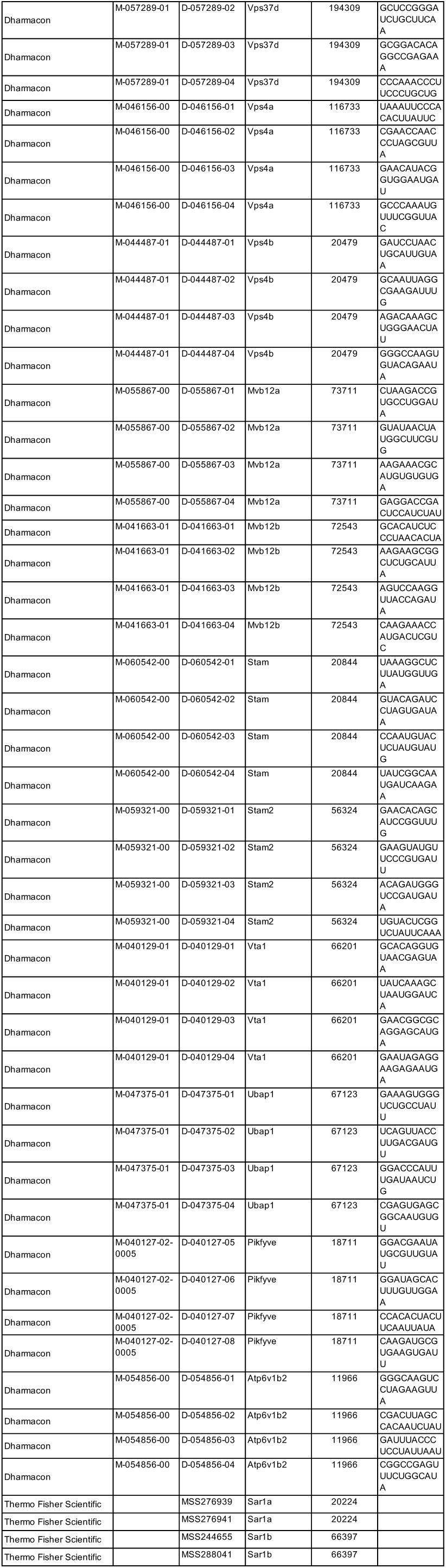

## References

1. Ishikawa, H. & Barber, G. N. STING is an endoplasmic reticulum adaptor that facilitates innate immune signalling. Nature 455, 674–678 (2008).

2. Zhong, B. et al. The adaptor protein MITA links virus-sensing receptors to IRF3 transcription factor activation. Immunity 29, 538–550 (2008).

3. Sun, W. et al. ERIS, an endoplasmic reticulum IFN stimulator, activates innate immune signaling through dimerization. Proc Natl Acad Sci U S A 106, 8653– 8658 (2009).

4. Jin, L. et al. MPYS, a novel membrane tetraspanner, is associated with major histocompatibility complex class II and mediates transduction of apoptotic signals. Mol Cell Biol 28, 5014–5026 (2008).

5. Barber, G. N. STING: infection, inflammation and cancer. Nat Rev Immunol 15, 760–770 (2015).

6. Motwani, M., Pesiridis, S. & Fitzgerald, K. A. DNA sensing by the cGAS-STING pathway in health and disease. Nat Rev Genet 20, 657–674 (2019).

7. Decout, A., Katz, J. D., Venkatraman, S. & Ablasser, A. The cGAS-STING pathway as a therapeutic target in inflammatory diseases. Nat Rev Immunol 21, 548–569 (2021).

8. Wu, J. et al. Cyclic GMP-AMP is an endogenous second messenger in innate immune signaling by cytosolic DNA. Science 339, 826–830 (2013).

9. Sun, L., Wu, J., Du, F., Chen, X. & Chen, Z. J. Cyclic GMP-AMP synthase is a cytosolic DNA sensor that activates the type I interferon pathway. Science 339, 786–791 (2013).

10. Mukai, K. et al. Activation of STING requires palmitoylation at the Golgi. Nat Commun 7, 11932 (2016).

11. Kemmoku, H., Kuchitsu, Y., Mukai, K. & Taguchi, T. Specific association of TBK1 with the trans-Golgi network following STING stimulation. Cell Struct Funct 47, 19–30 (2022).

12. Kemmoku, H. et al. Single-molecule localization microscopy reveals STING clustering at the trans-Golgi network through palmitoylation-dependent accumulation of cholesterol. Nat Commun 15, 220 (2024).

13. Kuchitsu, Y. et al. STING signalling is terminated through ESCRT-dependent microautophagy of vesicles originating from recycling endosomes. Nat Cell Biol 25, 453–466 (2023).

14. Kuchitsu, Y. & Taguchi, T. Lysosomal microautophagy: an emerging dimension in mammalian autophagy. Trends Cell Biol S0962–8924(23)00238 (2023).

15. Vietri, M., Radulovic, M. & Stenmark, H. The many functions of ESCRTs. Nat Rev Mol Cell Biol 21, 25–42 (2020).

16. Raiborg, C. & Stenmark, H. The ESCRT machinery in endosomal sorting of ubiquitylated membrane proteins. Nature 458, 445–452 (2009).

17. Taguchi, T. Membrane traffic governs the STING inflammatory signalling. J Biochem 174, 483–490 (2023).

18. Jefferies, H. B. et al. A selective PIKfyve inhibitor blocks PtdIns(3,5)P(2) production and disrupts endomembrane transport and retroviral budding. EMBO Rep 9, 164–170 (2008).

19. Hasegawa, J., Strunk, B. S. & Weisman, L. S. PI5P and PI(3,5)P_2_: Minor, but Essential Phosphoinositides. Cell Struct Funct 42, 49–60 (2017).

20. Cai, X. et al. PIKfyve, a class III PI kinase, is the target of the small molecular IL-12/IL-23 inhibitor apilimod and a player in Toll-like receptor signaling. Chem Biol 20, 912–921 (2013).

21. Choy, C. H. et al. Lysosome enlargement during inhibition of the lipid kinase PIKfyve proceeds through lysosome coalescence. J Cell Sci 131, jcs213587 (2018).

22. Gentili, M. et al. ESCRT-dependent STING degradation inhibits steady-state and cGAMP-induced signalling. Nat Commun 14, 611 (2023).

23. Balka, K. R. et al. Termination of STING responses is mediated via ESCRT-dependent degradation. EMBO J 42, e112712 (2023).

24. Raiborg, C. et al. FYVE and coiled-coil domains determine the specific localisation of Hrs to early endosomes. J Cell Sci 114, 2255–2263 (2001).

25. Bache, K. G., Brech, A., Mehlum, A. & Stenmark, H. Hrs regulates multivesicular body formation via ESCRT recruitment to endosomes. J Cell Biol 162, 435–442 (2003).

26. Edgar, J. R., Eden, E. R. & Futter, C. E. Hrs- and CD63-dependent competing mechanisms make different sized endosomal intraluminal vesicles. Traffic 15, 197–211 (2014).

27. Huber, S. T., Mostafavi, S., Mortensen, S. A. & Sachse, C. Structure and assembly of ESCRT-III helical Vps24 filaments. Sci Adv 6, eaba4897 (2020).

28. Rutherford, A. C. et al. The mammalian phosphatidylinositol 3-phosphate 5-kinase (PIKfyve) regulates endosome-to-TGN retrograde transport. J Cell Sci 119, 3944–3957 (2006).

29. Ikonomov, O. C. et al. The phosphoinositide kinase PIKfyve is vital in early embryonic development: preimplantation lethality of PIKfyve-/- embryos but normality of PIKfyve+/- mice. J Biol Chem 286, 13404–13413 (2011).

30. Takasuga, S. et al. Critical roles of type III phosphatidylinositol phosphate kinase in murine embryonic visceral endoderm and adult intestine. Proc Natl Acad Sci U S A 110, 1726–1731 (2013).

31. Rivero-Ríos, P. & Weisman, L. S. Roles of PIKfyve in multiple cellular pathways. Curr Opin Cell Biol 76, 102086 (2022).

32. Jin, N. et al. VAC14 nucleates a protein complex essential for the acute interconversion of PI3P and PI(3,5)P(2) in yeast and mouse. EMBO J 27, 3221– 3234 (2008).

33. Botelho, R. J., Efe, J. A., Teis, D. & Emr, S. D. Assembly of a Fab1 phosphoinositide kinase signaling complex requires the Fig4 phosphoinositide phosphatase. Mol Biol Cell 19, 4273–4286 (2008).

34. Barlow-Busch, I., Shaw, A. L. & Burke, J. E. PI4KA and PIKfyve: Essential phosphoinositide signaling enzymes involved in myriad human diseases. Curr Opin Cell Biol 83, 102207 (2023).

35. Li, S. et al. Mutations in PIP5K3 are associated with François-Neetens mouchetée fleck corneal dystrophy. Am J Hum Genet 77, 54–63 (2005).

36. Kotoulas, A. et al. A novel PIKFYVE mutation in fleck corneal dystrophy. Mol Vis 17, 2776–2781 (2011).

37. Gee, J. A. et al. Identification of novel PIKFYVE gene mutations associated with Fleck corneal dystrophy. Mol Vis 21, 1093–1100 (2015).

38. Mei, S. et al. Disruption of PIKFYVE causes congenital cataract in human and zebrafish. Elife 11, e71256 (2022).

39. Koga, D., Kusumi, S., Ushiki, T. & Watanabe, T. Integrative method for three-dimensional imaging of the entire Golgi apparatus by combining thiamine pyrophosphatase cytochemistry and array tomography using backscattered electron-mode scanning electron microscopy. Biomed Res 38, 285–296 (2017).

40. Bray, N. L., Pimentel, H., Melsted, P. & Pachter, L. Near-optimal probabilistic RNA-seq quantification. Nat Biotechnol 34, 525–527 (2016).

41. Hänzelmann, S., Castelo, R. & Guinney, J. GSVA: gene set variation analysis for microarray and RNA-seq data. BMC Bioinformatics 14, 7 (2013).

